# Number of Synergies Impacts Sensitivity of Gait to Weakness and Contracture

**DOI:** 10.1101/2021.06.06.447290

**Authors:** Elijah C. Kuska, Naser Mehrabi, Michael H. Schwartz, Katherine M. Steele

**Affiliations:** Department of Mechanical Engineering, University of Washington, Seattle, WA; Center for Gait & Motional Analysis, Gillette Children’s Specialty Healthcare, St. Paul, MN

**Keywords:** Altered motor modules, Impaired motor control, Musculoskeletal impairment, Walking simulation, Control Complexity

## Abstract

Muscle activity during gait can be described by a small set of synergies, weighted groups of muscles, that are often theorized to reflect underlying neural control. For people with neurologic injuries, like in cerebral palsy or stroke, even fewer *(e*.*g*., < 5) synergies are required to explain muscle activity during gait. This reduction in synergies is thought to reflect simplified control strategies and is associated with impairment severity and treatment outcomes. Individuals with neurologic injuries also develop secondary musculoskeletal impairments, like weakness or contracture, that can also impact gait. The combined impacts of simplified control and musculoskeletal impairments on gait remains unclear. In this study, we use a musculoskeletal model constrained to synergies to simulate unimpaired gait. We vary the number of synergies (3-5), while simulating muscle weakness and contracture to examine how altered control impacts sensitivity to muscle weakness and contracture. Our results highlight that reducing the number of synergies increases sensitivity to weakness and contracture. For example, simulations using five-synergy control tolerated 40% and 51% more knee extensor weakness than those using four- and three-synergy control, respectively. Furthermore, the model became increasingly sensitive to contracture and proximal muscle weakness, such as hamstring and hip flexor weakness, when constrained to four- and three-synergy control. However, the model’s sensitivity to weakness of the plantarflexors and smaller bi-articular muscles was not affected by the number of synergies. These findings provide insight into the interactions between altered control and musculoskeletal impairments, emphasizing the importance of incorporating both in future simulation studies.

## 1. Introduction

Muscle synergy analysis decomposes muscle excitations to identify common patterns of co-activation during dynamic activities. These patterns are theorized to reflect modular spinal and supraspinal networks that are used to control movement (Kargo et al., 2010; Stein, 2008) and reduce the dimensionality of neuromuscular control (Tresch and Jarc, 2009). More recently, these analyses have provided a tool to quantify the complexity of an individual’s motor control (Cappellini et al., 2016; Clark et al., 2010; Steele et al., 2015; Tang et al., 2015). Extending synergy analyses to clinical populations, like children with cerebral palsy (CP) and individuals post-stroke, has improved our understanding of neuromuscular impairments. For example, muscle activity during walking for children with CP and individuals post-stroke can be explained by fewer synergies than nondisabled peers, which has been associated with impaired function (Cheung et al., 2012) and treatment outcomes (Oudenhoven et al., 2017; Schwartz et al., 2016; Shuman et al., 2019a, 2018). These individuals also commonly develop secondary musculoskeletal impairments. In particular, muscle contracture and weakness are common (Gage et al., 2009; O’Dwyer et al., 1996). The interactions between altered control and secondary musculoskeletal impairments make identifying the causal mechanisms underlying gait pathologies challenging and limit our ability to effectively intervene.

Computational models of the neuromusculoskeletal system enable evaluation of hypotheses about relationships between impairment mechanisms. Previous studies used modeling and simulation to examine how weakness (van der Krogt et al., 2012), contracture (Fox et al., 2018), and the number of synergies (Mehrabi et al., 2019) may impact gait. Results highlighted that weakness, contracture, and reliance on fewer synergies, make achieving a typical gait pattern more difficult. However, these impairments were imposed in isolation; therefore, not addressing if altered control inhibits adaptation to musculoskeletal impairments (Fox et al., 2018). A previous simulation study examined the combined effects of these impairments in the presence of aberrant musculoskeletal geometries. Results highlighted that the number of synergies may influence how musculoskeletal impairments impact gait and suggested that altered muscle-tendon properties, rather than impaired motor control, were the primary cause of abnormal gait in a case study of a child with CP (Falisse et al., 2020).

The purpose of the present study was to examine the interactions between neuromuscular and musculoskeletal impairments on unimpaired gait. Specifically, we used musculoskeletal simulation to examine how the number of synergies controlling movement alters the sensitivity of unimpaired gait to weakness and contracture. We used a direct collocation framework (Mehrabi et al., 2019) to generate tracking simulations while varying the number of synergies and simulating progressive weakness and contracture. Reducing the number of synergies controlling movement restricts control space and flexibility, thus, we hypothesized that control strategies constrained to fewer synergies would (1) reduce the amount of weakness and contracture the simulation can tolerate before unimpaired gait is irreplicable and (2) exacerbate the increases in muscle activity required to replicate unimpaired gait with weakness and contracture.

## 2. Methods

### 2.1 Musculoskeletal model

We built a sagittal plane, musculoskeletal model based on the planar model of (Geyer and Herr, 2010) and added the rectus femoris similar to (Dorn et al., 2015) in MapleSim (Maplesoft, Inc). The model consisted of seven rigid body segments - one combined head, arms, and torso (HAT) and three segments per leg (thigh, shank, and foot) – linked by hinge joints (Geyer and Herr, 2010) (Figure 1). Eight Hill-type musculotendinous units per leg actuated the model’s nine kinematic degrees of freedom (DoF): biarticular hamstring (HAM), gluteus maximus (GLU), iliopsoas (IP), rectus femoris (RF), vasti (VAS), gastrocnemius (GAS), soleus (SOL), and tibialis anterior (TA) (Dorn et al., 2015; Mehrabi et al., 2019). Ten continuous coulomb friction contact spheres were placed equidistantly along each foot to simulate ground contact (Brown and McPhee, 2016). Sphere contact stiffness was hand-tuned to a value of 848500 N/m to prevent unrealistic foot movements and spikes in ground reaction forces (Mehrabi et al., 2019).

**Figure 1:**
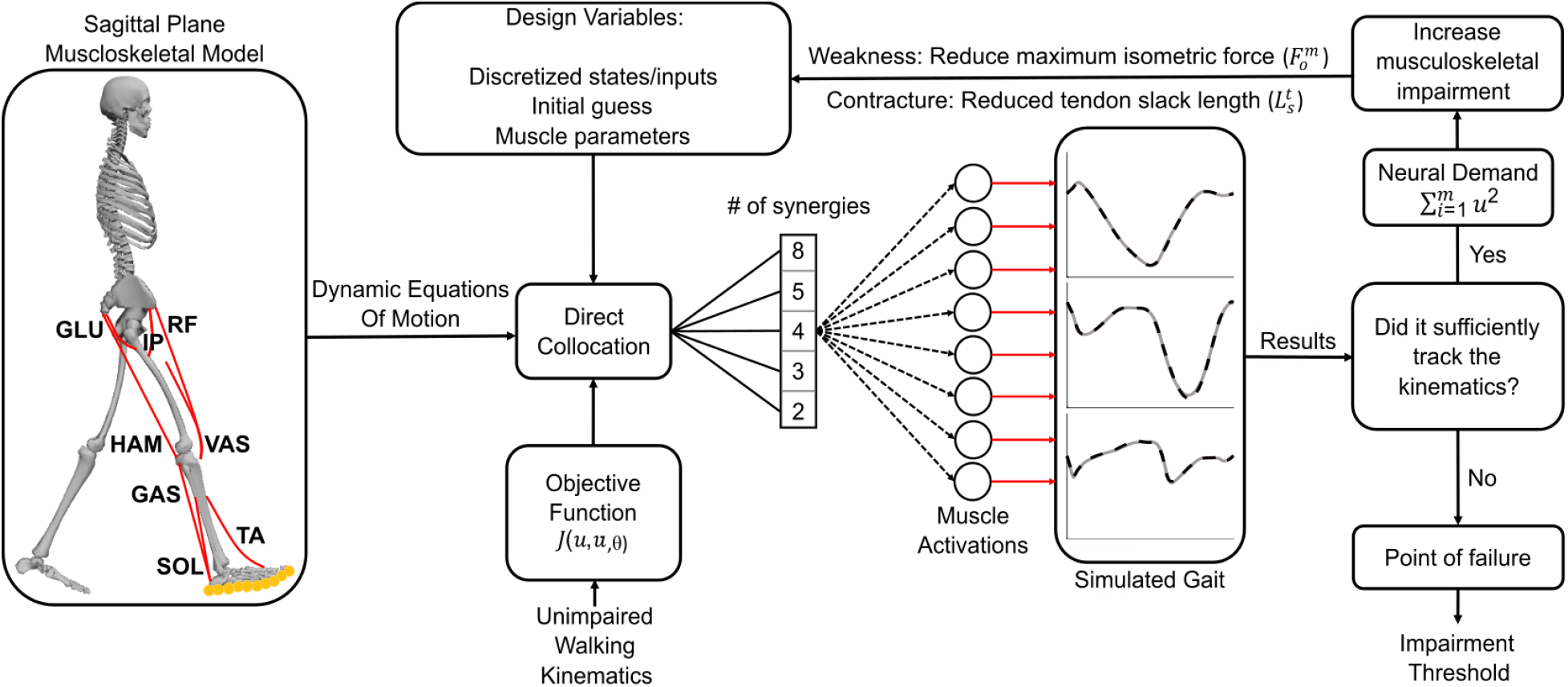
A two-dimensional sagittal plane musculoskeletal model and synergy simulation framework tracked unimpaired gait kinematics. The model had nine degrees of freedom, including right and left leg hip, knee, and ankle flexion, actuated by eight muscles per leg. Fixed sets of synergies constrained control, forcing the direct collocation algorithm to solve for synergy activations. The objective function minimized deviations from unimpaired kinematics and the sum of muscle activations (u) squared (neural demand). Weakness, simulated by a reduction in maximum isometric force 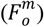, and contracture, simulated by a reduction in tendon slack length 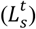, were progressively increased for each muscle or muscle group until the simulation failed to replicate unimpaired gait. Kinematic deviations and convergence determined the success of the simulation. The primary outcomes were (1) musculoskeletal impairment thresholds, defined by the amount of weakness or contracture before failure, and (2) neural demand of each gait cycle.

### 2.2 Optimization

Dynamic equations of motion were exported from MapleSim to a direct collocation (DC) optimal control framework (Figure 1) within MATLAB (Mathworks, Inc) (Mehrabi et al., 2019). DC is a trajectory optimization method that discretizes states along a path and solves for each discretization point simultaneously. DC has become popular because of its ability to accurately and rapidly simulate motion (Ackermann and van den Bogert, 2010; De Groote et al., 2016; Kinney et al., 2013).

Within MATLAB, ADiGator (Weinstein and Rao, 2017) converted the trajectory optimization into a nonlinear program which MATLAB’s interior-point optimizer (IPOPT) (Wächter and Biegler, 2006) solved. The framework generated tracking simulations that minimized deviations from desired kinematics, the amount of muscle activation required - which we termed “neural demand” (Ackermann and van den Bogert, 2010; Mehrabi et al., 2019) - and a smoothing term for neural control:

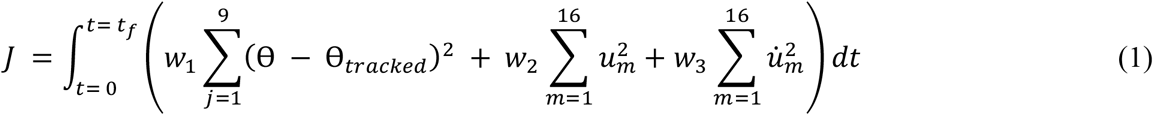

where ⊖ - ⊖_tracked_ represents deviations from tracked kinematics, and *u* represents muscle activations with 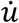 derivatives. Neural demand acted as an energetic estimator in lieu of a metabolic model as muscle activations require substantial metabolic energy (Lemaire et al., 2019), are a primary determinant of human energy expenditure (Umberger et al., 2003), and correlate with the metabolic cost of walking (Hortobágyi et al., 2011; Silder et al., 2012). Weighting factors (w_1_, w_2_, and w_3_) applied to the tracking, neural demand, and neural derivative terms were set to 5000, 35, and 0.05, respectively. These weights were previously used to simulate unimpaired gait (Mehrabi et al., 2019). Gait replication was the primary goal, thus tracking error was most heavily weighted. Average walking kinematics from a previous study of unimpaired individuals (Liu et al., 2008) were tracked (⊖_*tracked*_). To reduce computation time, symmetry was assumed (Ankarali et al., 2015). Preliminary simulations were initialized with a null guess: perfect state matching and zeroed controls, whereafter simulations used the preliminary solutions to initialize the optimization.

### 2.3 Neuromuscular Control

We implemented a neuromuscular controller within the DC framework (Figure 1) that varied the number of synergies controlling each leg (Steele et al., 2015). Non-negative matrix factorization (NNMF) (Lee and Seung, 1999) generated synergies from an initial simulation that controlled each muscle individually (*i*.*e*., not constrained by synergies). Briefly, NNMF decomposes control signals into weights and activations by a multiplicative update algorithm that minimizes deviations from the original signal. Thus, the chosen sets of 3-5 synergies represented the groups of muscles that would explain the greatest variance in muscle activity from the initial simulation.

The synergies constrained muscle activation patterns during tracking simulations, where fewer synergies were used to simulate more severe neuromuscular impairments (Cheung et al., 2012; Steele et al., 2015). We selected 3-5 synergies as this reflects the control complexities observed in CP (Bekius et al., 2020; Shuman et al., 2019a; Steele et al., 2015; Tang et al., 2015), individuals post-stroke (Clark et al., 2010), and nondisabled individuals (Rozumalski et al., 2017; Shuman et al., 2019b). Two synergy control, sometimes seen in stroke (Clark et al., 2010) and CP (Shuman et al., 2019a; Tang et al., 2015), was excluded because of its inability to track unimpaired gait (Mehrabi et al., 2019).

### 2.4 Musculoskeletal Impairments

We simulated weakness by reducing a muscle or muscle group’s maximum isometric force 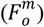 (Fox et al., 2018; Ong et al., 2019; Steele et al., 2012; van der Krogt et al., 2012). Each muscle was weakened individually, along with both plantarflexors (PFlex = GAS + SOL), knee extensors (KExt = RF + VAS), knee flexors (KFlex = GAS + HAM), hip extensors (HExt = GLU + HAM), and hip flexors (HFlex = IP + RF). To examine generalized weakness, all muscles were weakened simultaneously (ALL).

We altered the original model (Mehrabi et al., 2019) to permit tendon strain and simulated contracture by reducing a muscle or muscle group’s tendon slack length 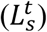 (Fox et al., 2018; Steele and Lee, 2014). We simulated contracture for SOL, GAS, HAM, and PFlex. These muscles exhibit contracture among children with CP (Barber et al., 2011; Handsfield et al., 2016; Wiley and Damiano, 1998) and stroke survivors (Diong and Herbert, 2015; Halar et al., 1978; Kwah et al., 2012), and are thought to contribute to pathologic gait (Graham et al., 2016; O’Dwyer et al., 1989; Ong et al., 2019; Steele and Lee, 2014).

### 2.5 Analyses

We progressively increased weakness or contracture, using 1% increments for weakness and 0.1% increments for contracture, until the simulation failed to replicate unimpaired gait. Points of failure, defined by either the average root-mean-squared error (RMSE) for a DoF surpassing one degree or the simulation’s inability to converge after 2500 iterations, were considered weakness and contracture thresholds. Thresholds demonstrate gait’s robustness to weakness or contracture. Differences in thresholds across controls highlight how reducing the number of synergies used by the control strategy can impact unimpaired gait’s sensitivity.

To further examine interactions between the neuromuscular and musculoskeletal impairments, we analyzed the neural demand required to replicate gait as weakness and contracture progressively increased with five-, four, and three-synergy control (Figure 4a). As noted above, neural demand was a term in the optimization’s objective function and calculated as the sum of squared activations. Neural demand from five-synergy control were subtracted from four- and three-synergy control, highlighting differences in neural demand from baseline control (Figure 4b). We fit quadratic curves to the neural demand differences to characterize the compounding effects of weakness and contracture on the amount of muscle activation required to track unimpaired gait. The second-order coefficient of a quadratic fit describes the steepness of the curvature. Greater second-order coefficients indicate larger increases and more rapid deviations in neural demand relative to five-synergy control (Figure 4b). To facilitate comparison, the second-order coefficients were normalized to the maximum observed for each impairment (*i*.*e*., weakness and contracture).

## 3. Results

### 3.1 Weakness

The tolerance to weakness decreased as the control strategy used fewer synergies to track unimpaired gait (Figure 2). When weakness was imposed, five-synergy control tolerated on average 27% and 35% more weakness than four- and three-synergy control, respectively. For generalized weakness (ALL), five-synergy control tolerated up to a 45% reduction in total body strength (*i*.*e*., the simulation failed to track unimpaired gait with five-synergies when all muscles were weakened by 45%). Four- and three-synergy control tolerated similar reductions in total body strength (44% and 45%, respectively), indicating the number of synergies had little to no effect on generalized weakness thresholds.

**Figure 2:**
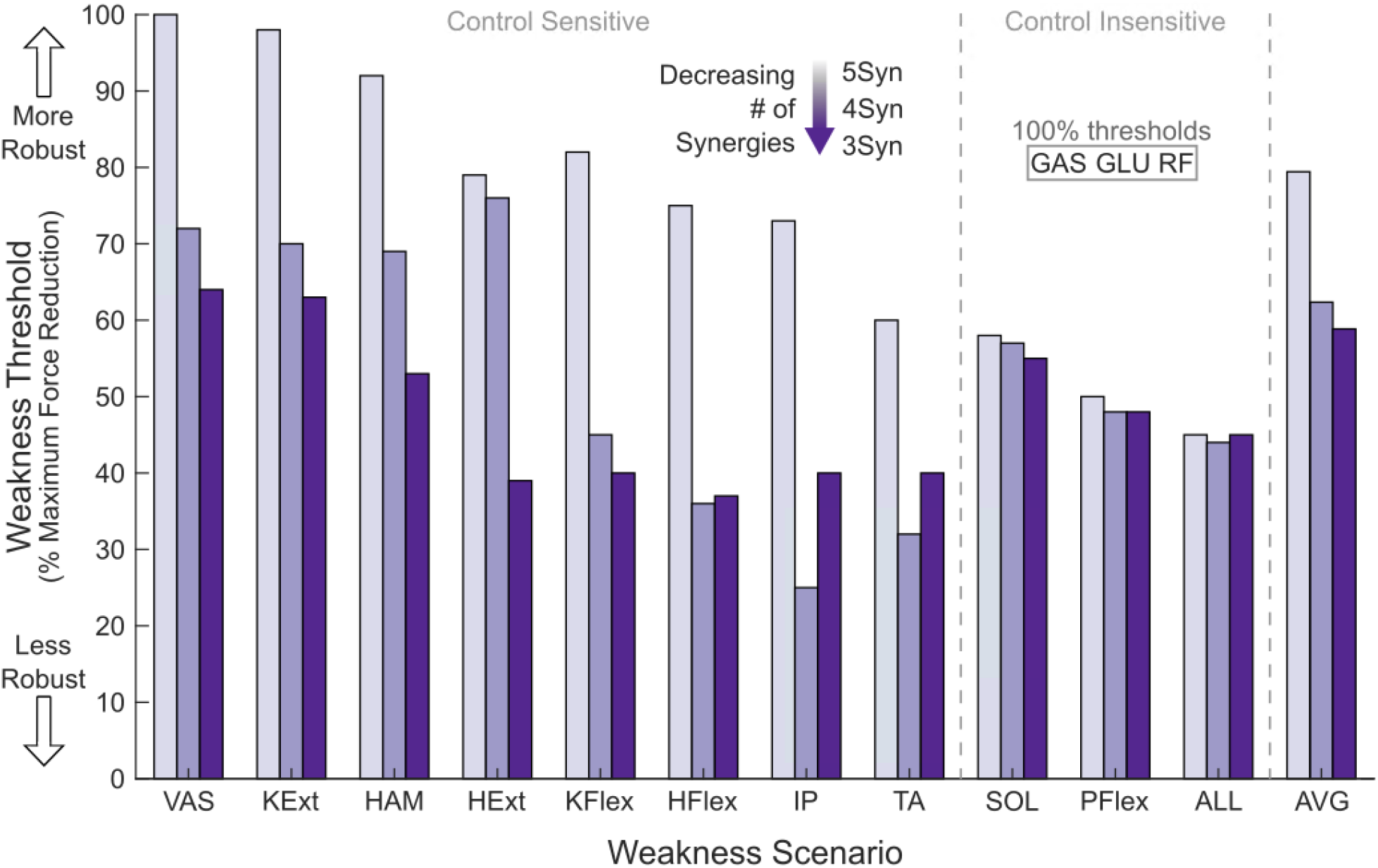
Weakness thresholds for five-, four-, and three-synergy control. A greater weakness threshold indicates muscle groups that are more robust to weakness while tracking unimpaired gait. Three muscles (GAS, GLU, and RF) had a weakness threshold of 100% (*i*.*e*., could be removed entirely from the simulation without impairing gait) for all synergy controls. Muscles were considered control sensitive or control insensitive based on the average threshold differences between five-, four-, and three-synergy control. If the average difference in weakness thresholds between five-, four-, and three-synergy control exceeded 7.7%, it was considered control sensitive. Control sensitivity indicated that changes in the number of synergies used by the control strategy altered weakness thresholds.

The weakness thresholds for specific muscle groups varied. We deemed muscle groups control sensitive if the average difference in weakness thresholds between five-, four-, and three-synergy control exceeded 7.7%. This value represents the standard deviation in lower limb muscle volume and strength for TD children (Handsfield et al., 2016; Knarr et al., 2013). Control sensitive muscle groups included the VAS, knee extensors (VAS + RF), HAM, hip extensors (HAM + GLU), knee flexors (GAS + HAM), hip flexors (IP + RF), IP, and TA (Figure 2). Within the control sensitive group, five-synergy control could replicate unimpaired gait with, on average, 34% and 41% more weakness than four- and three-synergy control, respectively. Within the control sensitive group, four- and three-synergy control were least robust to TA, IP, and hip flexor (IP + RF) weakness.

The muscle groups that were not sensitive to changes in the number of synergies (*i*.*e*., control insensitive) included the GAS, GLU, RF, SOL, and PFlex (SOL + GAS). Three muscles, the GAS, GLU, and RF, could be fully weakened (i.e., completely removed from the simulation with a weakness threshold of 100%) without preventing unimpaired gait for all synergy controls. The SOL and plantarflexors (SOL + GAS) had weakness thresholds of 58% and 50% for five-synergy control, respectively, with minimal differences (0-3 percentage points) in weakness thresholds with fewer synergies.

### 3.2 Contracture

The tolerance to contracture decreased as the control strategy used fewer synergies to track unimpaired gait (Figure 3). On average, five-synergy control tolerated 24.7% and 57.1% greater reductions in tendon slack length 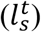 when compared to four- and three-synergy control. However, contracture thresholds varied between muscle groups. All three plantarflexor scenarios (SOL, GAS, SOL+GAS) showed similar robustness to contracture, with a 1.6 percentage point drop in contracture threshold from five-to four-synergy control, but then no further reduction in contracture threshold for three-synergy control. Similarly, HAM contracture had a 1.6 percentage point drop in threshold from five-to four-synergy control, but then had a 7.6 percentage point drop in threshold with three-synergy control.

**Figure 3:**
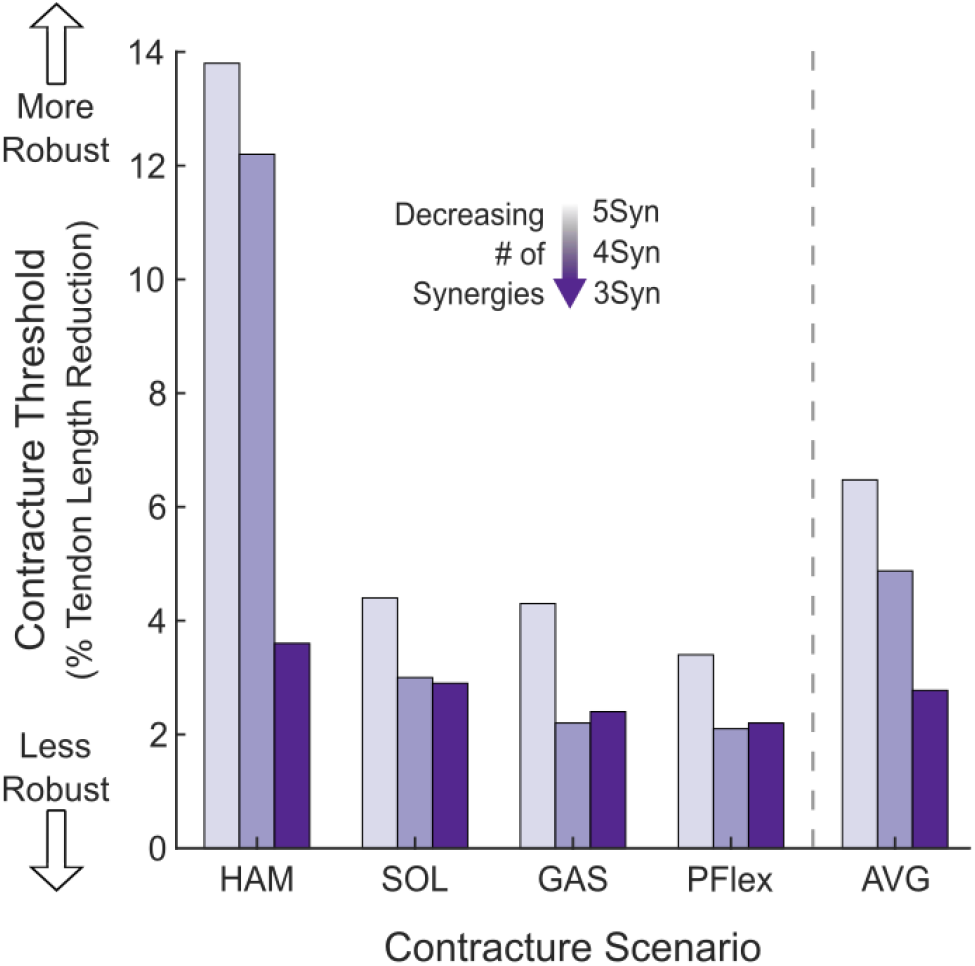
Contracture thresholds with five-, four-, and three-synergy control. A greater contracture threshold indicates muscle groups that are more robust to contracture while tracking unimpaired gait.

### 3.3 Neural Demand

Weakness, contracture, and the number of synergies used by the control strategy impacted the neural demand (sum of squared muscle activations) during gait. Five-synergy control’s neural demand was most sensitive to ALL, PFlex, SOL, and hip flexor (HFL and HFL + RF) weakness and least sensitive to knee extensor (RF, VAS, and VAS + RF), GLU, and GAS weakness. Contracture of the plantarflexors (SOL, PFlex, and GAS) increased five-synergy control’s neural demand more than HAM contracture. Without weakness or contracture, neural demand increased by 16.7% with four-synergy control compared to five-synergy control, and was further elevated by 3.7% with three-synergy control (Figure 4a, y-axis intercept).

**Figure 4:**
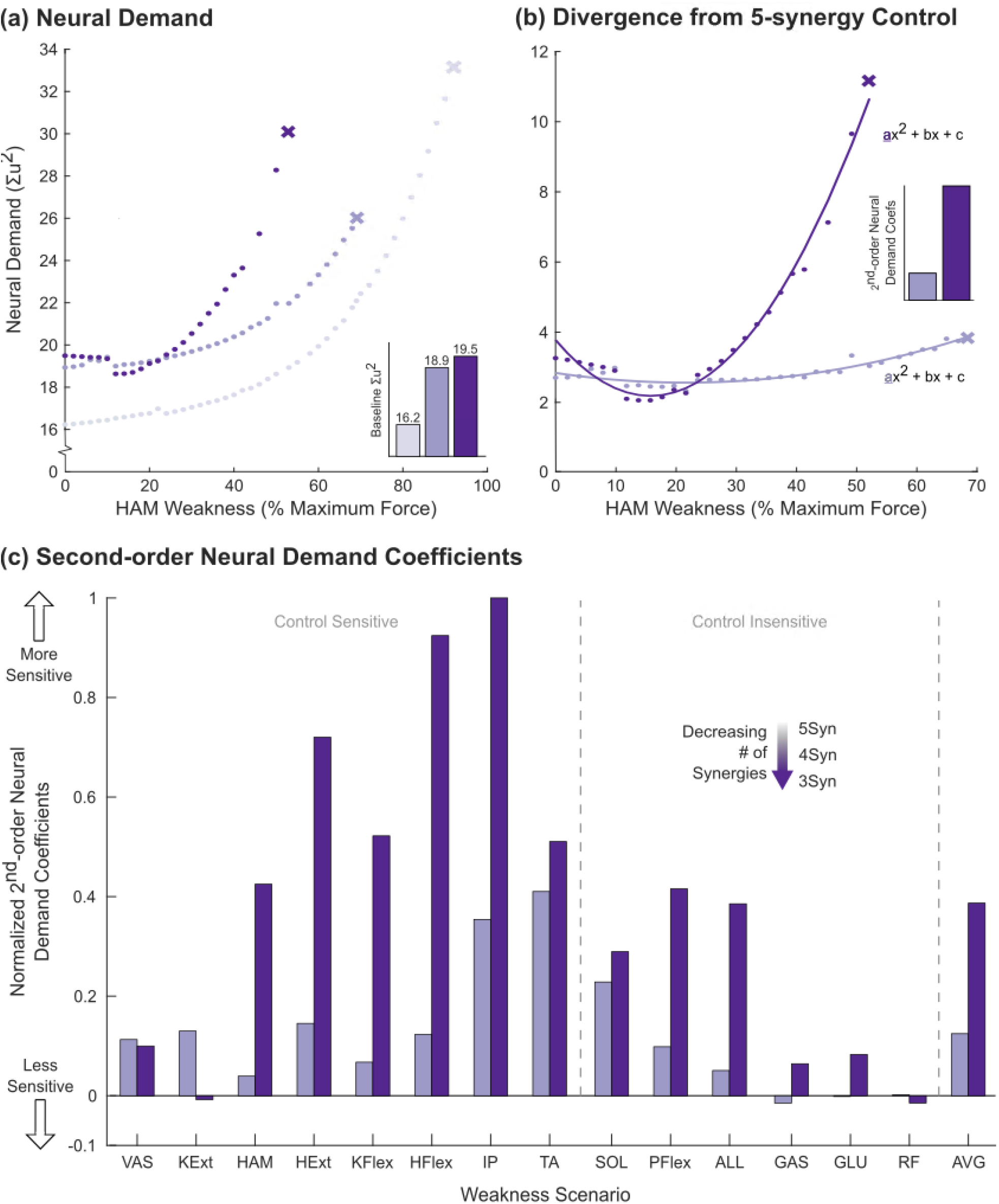
Neural demand – the summation of muscle activations squared - results for weakness simulations. (a) Neural demand values for five-, four-, and three-synergy control as hamstring (HAM) weakness, used as an example, was progressively increased. Weakness thresholds (X) indicate when simulations could no longer track unimpaired gait. Baseline neural demand (no weakness or contracture) was higher when control was constrained to fewer synergies (y-axis intercept). (b) Points represent five-synergy control’s neural demand subtracted from four- and three-synergy neural demand. Differences from five-synergy control’s neural demand were fit with quadratic polynomials where second-order coefficients indicate neural demand’s sensitivity to weakness (*i*.*e*., how, as weakness progressed, neural demand increased and deviated from five-synergy control when control strategies were constrained to use fewer synergies). (c) Weakness second-order neural demand coefficients, normalized to the maximum (0.02) of four- and three-synergy control. Greater second-order coefficients indicate greater sensitivity and larger increases in neural demand relative to five-synergy control with increasing weakness.

The sensitivity of neural demand to weakness increased as the control strategy used fewer synergies for most muscles (Figure 4c). The quadratic fit comparing the difference in neural demand between three- and five-synergy control was 3.5x that of four-synergy control. That is, as weakness progressed, the neural demand of three-synergy control increased and deviated from baseline control more rapidly than four-synergy control. For example, when ALL muscles were weakened, the second-order coefficient increased from 0.05 for four-synergy control to 0.39 for three-synergy control. Deviations from five-synergy control were greatest for the control sensitive muscles, especially weakness of the IP and TA. In contrast, the number of synergies were less influential on neural demand for the control insensitive muscles, with GAS, GLU, and RF second-order coefficients close to zero.

The sensitivity of neural demand to contracture increased as the control strategy used fewer synergies (Figure 5). When averaged across the contracture scenarios, the second-order coefficient was 2.5x larger for three-than four-synergy control. Deviations from five-synergy control, as measured by the average second-order coefficient between four- and three-synergy control, were largest for contracture of the plantarflexors (SOL, GAS, and SOL + GAS) and smallest for HAM contracture.

**Figure 5:**
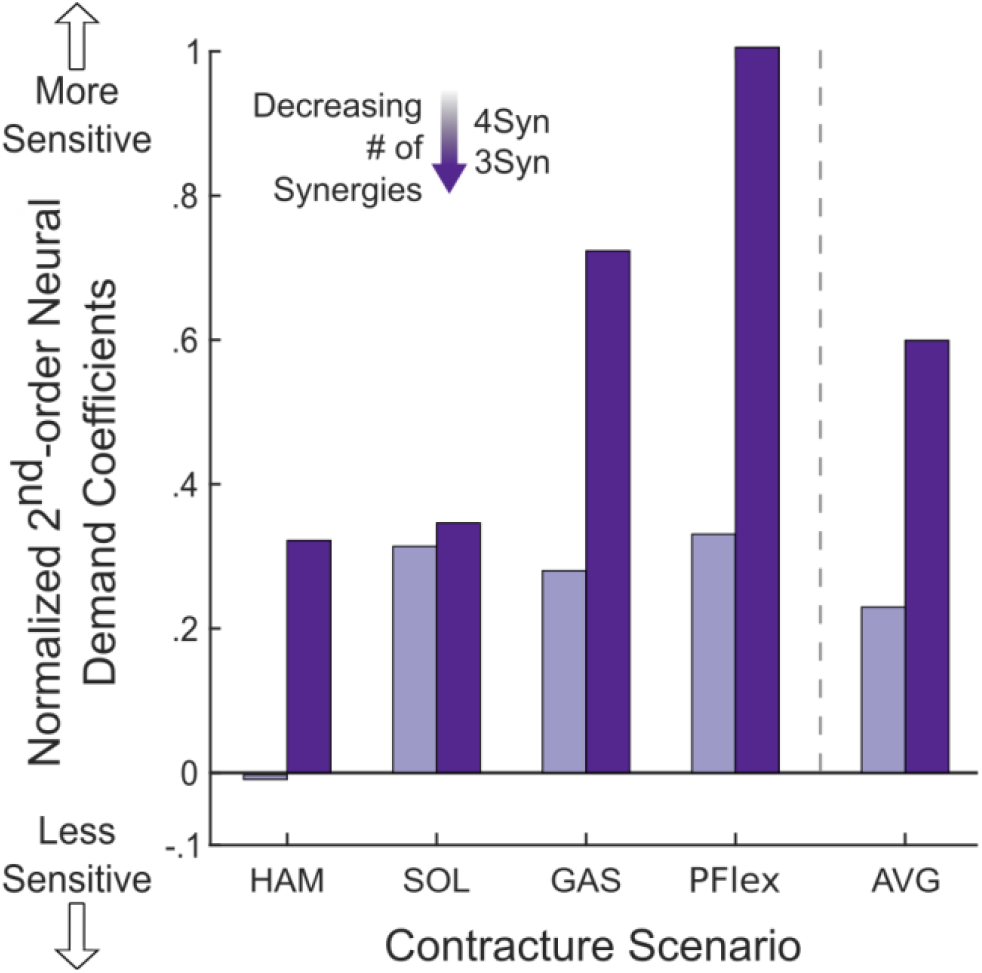
Contracture second-order neural demand – summation of muscle activations squared - coefficients, normalized to the maximum (1.46) of four- and three-synergy control. Greater second-order coefficients indicate increased sensitivity and larger increases in neural demand relative to five-synergy control as contracture progressed.

## 4. Discussion

This study found that impaired motor control (*e*.*g*., a control strategy constrained to fewer synergies) impacts how sensitive unimpaired gait is to musculoskeletal impairments. This supports both our hypotheses: control strategies constrained to use fewer synergies (1) tolerated less weakness and contracture while replicating unimpaired gait and (2) exacerbated the increases in muscle activation required to replicate unimpaired gait with weakness and contracture. Prior analyses have excluded altered control from systematic analyses on impacts of weakness or contracture on gait (Fox et al., 2018; Jason J Kutch and Valero-Cuevas, 2011; van der Krogt et al., 2012). Our results highlight interactions between altered control and musculoskeletal impairments which influence the sensitivity of gait and should be factored into future analyses, especially for populations with motor impairments.

The majority of muscles and muscle groups were sensitive to changes in the number of synergies used to track unimpaired gait. In general, weakness of proximal muscles and contracture of the hamstring and plantarflexors were most sensitive to changes in the number of synergies. Previous studies examining muscle weakness found that unimpaired gait was generally robust to weakness when control was not constrained to synergies (Fox et al., 2018; van der Krogt et al., 2012), but could only tolerate up to 3% reductions in tendon slack length (Fox et al., 2018). Our findings indicate similar trends for baseline control: five-synergy control was robust to the removal of several muscles and tolerated up to 3% reductions in tendon slack length for the plantarflexors. However, with four- and three-synergy control, simulations were less robust to weakness and contracture. Thus, the ability for the simulation to track unimpaired gait with weakness and contracture quickly diminished as the control strategy was constrained to use fewer synergies.

Our sagittal-plane simulations of unimpaired gait were most robust to the weakness of redundant and bi-articular muscles. Independent of the number of synergies, our simulations could track unimpaired gait with the removal of the gluteus maximus, rectus femoris, and gastrocnemius, which may be surprising due to the low redundancy measure (# of muscles divided # DoFs+1 (Sohn et al., 2019)) of our model compared to previous studies: 1.5 vs. 3.8 (van der Krogt et al., 2012) and 4.8 (Fox et al., 2018). Gluteus maximus results may be the most surprising because of the gluteus maximus’ ability to generate hip extensor moments (Arnold et al., 2005). However, our model and other previous studies found that redundant hip and knee extensors could actively compensate for gluteus maximus weakness (van der Krogt et al., 2012). Robustness to rectus femoris and gastrocnemius were previously reported (van der Krogt et al., 2012) and stem from lower maximum isometric forces and reduced abilities to accelerate joints per unit force (Arnold et al., 2005), allowing alternative control strategies to replicate unimpaired gait with as few as three synergies. In contrast, (Jason J. Kutch and Valero-Cuevas, 2011) found that removing the gastrocnemius and rectus femoris had the greatest impact on endpoint force generation, emphasizing a lack of redundancy in the musculoskeletal system . However, comparing this to our results from gait highlights how redundancy can be task-specific.

From a clinical perspective, it is surprising our results indicate that gluteus maximus, rectus femoris, and gastrocnemius weakness would not prevent unimpaired gait as these are common suspects of gait deviations in CP (Falisse et al., 2020; Ong et al., 2019; Steele et al., 2012). Model simplifications are likely not the sole cause as prior studies using more complex models found similar robustness to GAS, GLU, and RF weakness (van der Krogt et al., 2012). Our investigation and prior studies analyzed the impact of weakness during only unimpaired gait. GAS, GLU, and RF weakness may be more substantial in gait patterns that rely more on flexors, such as crouch gait in CP (Steele et al., 2012). Additionally, children with CP can develop bony deformities (Graham et al., 2016) and often have deficits in multiple muscle groups (Handsfield et al., 2016; Wiley and Damiano, 1998), likely affecting sensitivity to altered muscle-tendon properties.

The neural demand of unimpaired gait was most sensitive to total body and plantarflexor weakness and plantarflexor contracture, aligning with previous results indicating that impairments in these muscle groups lead to large increases in neural demand (van der Krogt et al., 2012). In addition, our study and previous findings highlight that knee extensor, gastrocnemius, gluteus maximus, and rectus femoris weakness have little to no effect on the neural demand of unimpaired gait (van der Krogt et al., 2012). Furthermore, the increases in neural demand required to adapt to weakness and contracture were exacerbated by fewer synergies. The magnitude of the exacerbating effect was largest for the weakness and contracture scenarios four- and three-synergy control were least robust to. These findings indicate that an individual’s muscle activation increases most when adapting to the to the musculoskeletal impairments they tolerate the least.

It is important to note that there are several limitations of this study. Simulations were a simplified representation of human gait that had simplified muscle paths and neglected active stability and control in the mediolateral direction. The addition of the mediolateral direction would likely highlight how unimpaired gait was sensitive to hip abductor weakness (van der Krogt et al., 2012) and simplified hip anatomy may have led to greater impairment thresholds for the hamstrings. Nonetheless, our simplified simulations found results similar to previous three-dimensional analyses of unimpaired gait’s robustness to weakness and contracture (Fox et al., 2018; van der Krogt et al., 2012) and highlights how, even in a simple model, fewer synergies can influence sensitivity to weakness and contracture. We also assumed bilateral symmetry which can be a poor assumption for individuals with hemiparesis. Future studies should examine unilateral muscle dysfunction and fewer synergies. Lastly, we only examined unimpaired gait and altered gait patterns may have different responses to the combined effects of fewer synergies, weakness, and contracture (Latash and Greg Anson, 1996). Future studies should apply similar methods to altered gait patterns as some gait deviations may be advantageous because they increase robustness to specific impairments.

This study investigated the interactions between neuromuscular and musculoskeletal impairments on gait. We found that when the control strategy was constrained to use fewer synergies, the likelihood of achieving an unimpaired gait pattern with weakness and contracture was further reduced. These results could be used to develop hypotheses for designing future interventions. For example, targeting plantarflexor weakness may be advantageous for an individual who uses fewer synergies because of its control insensitivity and importance in gait (Hall et al., 2011; Liu et al., 2008; van der Krogt et al., 2012). In conclusion, the severity with which musculoskeletal impairments influence gait is affected by the number of synergies required to explain one’s muscle activity during gait. Incorporating these factors into patient-specific models could improve our understanding of function for individuals with neuromuscular impairments.

## 5. Declaration of Competing Interest

There are no conflicts of interest to report.

## 6. Acknowledgements

This research was supported by the National Institute of Neurological Disorders and Stroke (NINDS) under grant number R01NS091056 in collaboration with Gillette Children’s Specialty Healthcare.

## Notes

### Competing Interest Statement

The authors have declared no competing interest.

### Summary of Updates

The figures and captions were moved to their appropriate position in the text to improve readability.

## References

Ackermann, M., van den Bogert, A.J., 2010. Optimality principles for model-based prediction of human gait. J. Biomech. 43, 1055–1060. https://doi.org/10.1016/j.jbiomech.2009.12.012

Ankarali, M.M., Sefati, S., Madhav, M.S., Long, A., Bastian, A.J., Cowan, N.J., 2015. Walking dynamics are symmetric (enough). J. R. Soc. Interface 12. https://doi.org/10.1098/rsif.2015.0209

Arnold, A.S., Anderson, F.C., Pandy, M.G., Delp, S.L., 2005. Muscular contributions to hip and knee extension during the single limb stance phase of normal gait: A framework for investigating the causes of crouch gait. J. Biomech. 38, 2181–2189. https://doi.org/10.1016/j.jbiomech.2004.09.036

Barber, L., Barrett, R., Lichtwark, G., 2011. Passive muscle mechanical properties of the medial gastrocnemius in young adults with spastic cerebral palsy. J. Biomech. 44, 2496–2500. https://doi.org/10.1016/j.jbiomech.2011.06.008

Bekius, A., Bach, M.M., van der Krogt, M.M., de Vries, R., Buizer, A.I., Dominici, N., 2020. Muscle Synergies During Walking in Children With Cerebral Palsy: A Systematic Review. Front. Physiol. 11, 632. https://doi.org/10.3389/fphys.2020.00632

Brown, P., McPhee, J., 2016. A Continuous Velocity-Based Friction Model for Dynamics and Control with Physically Meaningful Parameters. J. Comput. Nonlinear Dyn. 11. https://doi.org/10.1115/1.4033658

Cappellini, G., Ivanenko, Y.P., Martino, G., MacLellan, M.J., Sacco, A., Morelli, D., Lacquaniti, F., 2016. Immature spinal locomotor output in children with cerebral palsy. Front. Physiol. 7, 478. https://doi.org/10.3389/fphys.2016.00478

Cheung, V.C.K., Turolla, A., Agostini, M., Silvoni, S., Bennis, C., Kasi, P., Paganoni, S., Bonato, P., Bizzi, E., 2012. Muscle synergy patterns as physiological markers of motor cortical damage. Proc. Natl. Acad. Sci. U. S. A. 109, 14652–14656. https://doi.org/10.1073/pnas.1212056109

Clark, D.J., Ting, L.H., Zajac, F.E., Neptune, R.R., Kautz, S.A., 2010. Merging of healthy motor modules predicts reduced locomotor performance and muscle coordination complexity post-stroke. J. Neurophysiol. 103, 844–857. PMCID: PMC2822696. https://doi.org/10.1152/jn.00825.2009

De Groote, F., Kinney, A.L., Rao, A. V, Fregly, B.J., 2016. Evaluation of direct collocation optimal control problem formulations for solving the muscle redundancy problem. Ann. Biomed. Eng. 44, 2922–2936.

Diong, J., Herbert, R.D., 2015. Is ankle contracture after stroke due to abnormal intermuscular force transmission? J. Appl. Biomech. 31, 13–18. https://doi.org/10.1123/JAB.2014-0064

Dorn, T.W., Wang, J.M., Hicks, J.L., Delp, S.L., 2015. Predictive simulation generates human adaptations during loaded and inclined walking. PLoS One 10. https://doi.org/10.1371/journal.pone.0121407

Falisse, A., Pitto, L., Kainz, H., Hoang, H., Wesseling, M., Van Rossom, S., Papageorgiou, E., Bar-On, L., Hallemans, A., Desloovere, K., Molenaers, G., Van Campenhout, A., De Groote, F., Jonkers, I., 2020. Physics-Based Simulations to Predict the Differential Effects of Motor Control and Musculoskeletal Deficits on Gait Dysfunction in Cerebral Palsy: A Retrospective Case Study. Front. Hum. Neurosci. 14. https://doi.org/10.3389/fnhum.2020.00040

Fox, A.S., Carty, C.P., Modenese, L., Barber, L.A., Lichtwark, G.A., 2018. Simulating the effect of muscle weakness and contracture on neuromuscular control of normal gait in children. Gait Posture 61, 169–175. https://doi.org/10.1016/j.gaitpost.2018.01.010

Gage, J.R., Schwartz, M.H., Koop, S.E., Novacheck, T.F., 2009. The Identification and Treament of Gait Problems in Cerebral Palsy, Clinis in Developmental Medicine.

Geyer, H., Herr, H., 2010. A Muscle-reflex model that encodes principles of legged mechanics produces human walking dynamics and muscle activities. IEEE Trans. Neural Syst. Rehabil. Eng. 18, 263–273. https://doi.org/10.1109/TNSRE.2010.2047592

Graham, H.K., Rosenbaum, P., Paneth, N., Dan, B., Lin, J.P., Damiano, Di.L., Becher, J.G., Gaebler-Spira, D., Colver, A., Reddihough, Di.S., Crompton, K.E., Lieber, R.L., 2016. Cerebral palsy. Nat. Rev. Dis. Prim. 2. https://doi.org/10.1038/nrdp.2015.82

Halar, E.M., Stolov, W.C., Venkatesh, B., Brozovich, F. V., Harley, J.D., 1978. Gastrocnemius muscle belly and tendon length in stroke patients and able-bodied persons. Arch. Phys. Med. Rehabil. 59, 476–484.

Hall, A.L., Peterson, C.L., Kautz, S.A., Neptune, R.R., 2011. Relationships between muscle contributions to walking subtasks and functional walking status in persons with post-stroke hemiparesis. Clin. Biomech. (Bristol, Avon) 26, 509–515. https://doi.org/10.1016/j.clinbiomech.2010.12.010

Handsfield, G.G., Meyer, C.H., Abel, M.F., Blemker, S.S., 2016. Heterogeneity of muscle sizes in the lower limbs of children with cerebral palsy. Muscle and Nerve 53, 933–945. https://doi.org/10.1002/mus.24972

Hortobágyi, T., Finch, A., Solnik, S., Rider, P., DeVita, P., 2011. Association between muscle activation and metabolic cost of walking in young and old adults. J Gerontol A Biol Sci Med Sci 66, 541–547. https://doi.org/10.1093/gerona/glr008

Kargo, W.J., Ramakrishnan, A., Hart, C.B., Rome, L.C., Giszter, S.F., 2010. A simple experimentally based model using proprioceptive regulation of motor primitives captures adjusted trajectory formation in spinal frogs. J. Neurophysiol. 103, 573–590. https://doi.org/10.1152/jn.01054.2007

Kinney, A.L., Besier, T.F., D’Lima, D.D., Fregly, B.J., 2013. Update on grand challenge competition to predict in vivo knee loads. J. Biomech. Eng. 135, 21012. https://doi.org/10.1115/1.4023255

Knarr, B.A., Ramsay, J.W., Buchanan, T.S., Higginson, J.S., Binder-Macleod, S.A., 2013. Muscle volume as a predictor of maximum force generating ability in the plantar flexors post-stroke. Muscle Nerve 48, 971–976. https://doi.org/10.1002/mus.23835

Kutch, Jason J, Valero-Cuevas, F.J., 2011. Muscle redundancy does not imply robustness to muscle dysfunction. https://doi.org/10.1016/j.jbiomech.2011.02.014

Kutch, Jason J., Valero-Cuevas, F.J., 2011. Muscle redundancy does not imply robustness to muscle dysfunction. J. Biomech. 44, 1264–1270. https://doi.org/10.1016/j.jbiomech.2011.02.014

Kwah, L.K., Harvey, L.A., Diong, J.H.L., Herbert, R.D., 2012. Half of the adults who present to hospital with stroke develop at least one contracture within six months: An observational study. J. Physiother. 58, 41–47. https://doi.org/10.1016/S1836-9553(12)70071-1

Latash, M.L., Greg Anson, J., 1996. What are “normal movements” in atypical populations? Behav. Brain Sci. 19. https://doi.org/10.1017/s0140525x00041467

Lee, D.D., Seung, H.S., 1999. Learning the parts of objects by non-negative matrix factorization. Nature 401, 788–791. https://doi.org/10.1038/44565

Lemaire, K.K., Jaspers, R.T., Kistemaker, D.A., Soest, A.J.K. Van, Laarse, W.J.V. Der, 2019. Metabolic cost of activation and mechanical efficiency of mouse soleus muscle fiber bundles during repetitive concentric and eccentric contractions. Front. Physiol. 10. https://doi.org/10.3389/fphys.2019.00760

Liu, M.Q., Anderson, F.C., Schwartz, M.H., Delp, S.L., 2008. Muscle contributions to support and progression over a range of walking speeds. J Biomech 41, 3243–3252. https://doi.org/10.1016/j.jbiomech.2008.07.031

Mehrabi, N., Schwartz, M.H., Steele, K.M., 2019. Can altered muscle synergies control unimpaired gait? J. Biomech. 90, 84–91. https://doi.org/10.1016/j.jbiomech.2019.04.038

O’Dwyer, N.J., Ada, L., Neilson, P.D., 1996. Spasticity and muscle contracture following stroke. Brain 119, 1737–1749. https://doi.org/10.1093/brain/119.5.1737

O’Dwyer, N.J., Neilson, P.D., Nash, J., 1989. Mechanisms of muscle growth related to muscle contracture in cerebral palsy. Dev. Med. Child Neurol. https://doi.org/10.1111/j.1469-8749.1989.tb04034.x

Ong, C.F., Geijtenbeek, T., Hicks, J.L., Delp, S.L., 2019. Predicting gait adaptations due to ankle plantarflexor muscle weakness and contracture using physics-based musculoskeletal simulations. PLoS Comput. Biol. 15, e1006993. https://doi.org/10.1371/journal.pcbi.1006993

Oudenhoven, L., Marianna, R., Annet, D., Harlaar, J., van der Krogt, M., Buizer, A., 2017. Factors associated with long-term improvement after SDR surgery in children with spastic diplegia. Gait Posture 57, 272–273. https://doi.org/10.1016/j.gaitpost.2017.06.412

Rozumalski, A., Steele, K.M., Schwartz, M.H., 2017. Muscle synergies are similar when typically developing children walk on a treadmill at different speeds and slopes. J. Biomech. 64, 112–119. https://doi.org/10.1016/j.jbiomech.2017.09.002

Schwartz, M.H., Rozumalski, A., Steele, K.M., 2016. Dynamic motor control is associated with treatment outcomes for children with cerebral palsy. Dev. Med. Child Neurol. 58, 1139– 1145. https://doi.org/10.1111/dmcn.13126

Shuman, B.R., Goudriaan, M., Desloovere, K., Schwartz, M.H., Steele, K.M., 2019a. Muscle synergies demonstrate only minimal changes after treatment in cerebral palsy. J. Neuroeng. Rehabil. 16, 1–10. https://doi.org/10.1186/s12984-019-0502-3

Shuman, B.R., Goudriaan, M., Desloovere, K., Schwartz, M.H., Steele, K.M., 2019b. Muscle synergy constraints do not improve estimates of muscle activity from static optimization during gait for unimpaired children or children with cerebral palsy. Front. Neurorobot. 13, 1–17. https://doi.org/10.3389/fnbot.2019.00102

Shuman, B.R., Goudriaan, M., Desloovere, K., Schwartz, M.H., Steele, K.M., 2018. Associations Between Muscle Synergies and Treatment Outcomes in Cerebral Palsy Are Robust Across Clinical Centers. Arch. Phys. Med. Rehabil. 99, 2175–2182. https://doi.org/10.1016/j.apmr.2018.03.006

Silder, A., Besier, T., Delp, S.L., 2012. Predicting the metabolic cost of incline walking from muscle activity and walking mechanics. J. Biomech. 45, 1842–1849. https://doi.org/10.1016/j.jbiomech.2012.03.032

Sohn, M.H., Smith, D.M., Ting, L.H., 2019. Effects of kinematic complexity and number of muscles on musculoskeletal model robustness to muscle dysfunction. PLoS One 14, e0219779. https://doi.org/10.1371/journal.pone.0219779

Steele, K.M., Lee, S., 2014. Using dynamic musculoskeletal simulation to evaluate altered muscle properties in cerebral palsy, in: ASME 2014 Dynamic Systems and Control Conference, DSCC 2014. American Society of Mechanical Engineers. https://doi.org/10.1115/DSCC2014-5955

Steele, K.M., Rozumalski, A., Schwartz, M.H., 2015. Muscle synergies and complexity of neuromuscular control during gait in cerebral palsy. Dev. Med. Child Neurol. 57, 1176– 1182. https://doi.org/10.1111/dmcn.12826

Steele, K.M., van der Krogt, M.M., Schwartz, M.H., Delp, S.L., 2012. How much muscle strength is required to walk in a crouch gait? J. Biomech. 45, 2564–2569. https://doi.org/10.1016/j.jbiomech.2012.07.028

Stein, P.S.G., 2008. Motor pattern deletions and modular organization of turtle spinal cord. Brain Res. Rev. https://doi.org/10.1016/j.brainresrev.2007.07.008

Tang, L., Li, F., Cao, S., Zhang, X., Wu, D., Chen, X., 2015. Muscle synergy analysis in children with cerebral palsy. J. Neural Eng. 12. https://doi.org/10.1088/1741-2560/12/4/046017

Tresch, M.C., Jarc, A., 2009. The case for and against muscle synergies. Curr. Opin. Neurobiol. https://doi.org/10.1016/j.conb.2009.09.002

Umberger, B.R., Gerritsen, K.G., Martin, P.E., 2003. A model of human muscle energy expenditure. Comput. Methods Biomech. Biomed. Engin. 6, 99–111. https://doi.org/10.1080/1025584031000091678

van der Krogt, M.M., Delp, S.L., Schwartz, M.H., 2012. How robust is human gait to muscle weakness? Gait Posture 36, 113–119. https://doi.org/10.1016/j.gaitpost.2012.01.017

Wächter, A., Biegler, L.T., 2006. On the implementation of an interior-point filter line-search algorithm for large-scale nonlinear programming. Math. Program. 106, 25–57. https://doi.org/10.1007/s10107-004-0559-y

Weinstein, M.J., Rao, A. V., 2017. Algorithm 984: ADiGator, a toolbox for the algorithmic differentiation of mathematical functions in MATLAB using source transformation via operator overloading. ACM Trans. Math. Softw. 44. https://doi.org/10.1145/3104990

Wiley, M.E., Damiano, D.L., 1998. Lower-extremity strength profiles in spastic cerebral palsy. Dev. Med. Child Neurol. https://doi.org/10.1111/j.1469-8749.1998.tb15369.x

